# Finding reliable phenotypes and detecting artefacts among *in vivo* and *in vitro* assays to characterize the refractory transcriptional activator Sxy (TfoX) in *Escherichia coli*

**DOI:** 10.1101/284836

**Authors:** Ebtihal Y. Alshabib, Stephen F. Fitzgerald, Tzu-Chiao Chao, Rhiannon C. Cameron, Rosemary J. Redfield, Sunita Sinha, Andrew D.S. Cameron

**Affiliations:** Department of Biology, University of Regina, Regina, Saskatchewan, Canada; Department of Zoology, University of British Columbia, Vancouver, British Columbia, Canada

## Abstract

The Sxy (TfoX) protein is required for expression of a distinct subset of the genes regulated by the cAMP receptor protein (CRP) in the model organisms *Escherichia coli*, *Haemophilus influenzae*, and *Vibrio cholerae*. Genetic studies have established that CRP and Sxy co-activate transcription at gene promoters containing DNA binding sites called CRP-S sites. In contrast, CRP acts without Sxy at gene promoters containing canonical CRP-N sites, suggesting that Sxy makes physical contacts with CRP and/or DNA to assist in transcriptional activation at CRP-S promoters. Despite growing interest in Sxy’s activity as a transcription factor, Sxy remains poorly characterized due to a lack of reliable phenotypes in *E. coli*. Experiments are further hampered by growth inhibition and formation of inclusion bodies when Sxy is overexpressed. In this study we applied diverse phenotypic and molecular assays to test for postulated Sxy functions and interactions. Mutations in conserved regions of Sxy and truncations in the Sxy C-terminus abolish transcriptional activation of a CRP-S promoter, and a 37 amino acid truncation of the C-terminus relieves the growth inhibition normally caused by Sxy overexpression. Sxy was unable to augment weakened CRP interactions to restore carbon metabolism phenotypes. Bandshift analysis and chromatin pull-down assays of Sxy-CRP-DNA interactions yielded intriguing evidence of CRP-Sxy and Sxy-DNA physical interactions. However, despite the careful application of standard protein purification protocols and quality control steps for nickel affinity column purification, protein mass spectrometry revealed the enrichment of additional DNA-binding proteins in nickel column eluates, presenting a probable source of artefactual protein-protein and protein-DNA interaction results. These findings highlight the importance of extensive controls and phenotypic assays for the study of poorly characterized and recalcitrant proteins like Sxy.

## Introduction

Natural competence is the ability of bacteria to actively take up DNA from their environment. The core mechanisms that regulate natural competence are conserved in the model organisms *Haemophilus influenzae*, *Vibrio cholerae*, and *Escherichia coli* [1, 2]. In these species, competence is induced by two positive regulators, CRP (cAMP receptor protein, also called catabolite activator protein) and Sxy (also known as TfoX). CRP is the master regulator of a carbon-energy starvation response [3], while Sxy appears restricted to regulation of genes required for natural competence, DNA replication, and nucleotide metabolism [2,4–18].

CRP is a global regulator that activates expression of hundreds of genes in *E. coli* and other Gamma-proteobacteria [1]. Binding to the allosteric effector, cAMP, causes conformational changes that activate CRP as a DNA binding protein; CRP can then form protein-protein contacts with RNA polymerase and recruit the polymerase to nearby gene promoters [19–21]. CRP binds preferentially to DNA sequences described by the consensus half-site 5′-A_1_A_2_A_3_T_4_G_5_T_6_G_7_A_8_T_9_C_10_T_11_. We refer to these canonical CRP binding sites as CRP-N sites to distinguish them from non-canonical “CRP-S” sites characterized by C_6_ instead of T_6_ [1, 22]. Unlike gene promoters containing CRP-N sites, gene promoters with CRP-S sites require both CRP and Sxy for transcriptional activation. We previously quantified how both *E. coli* CRP (*Ec*CRP) and *H. influenzae* CRP (*Hi*CRP) demonstrate strong preferences for CRP-N over CRP-S sites [22], consistent with the finding that *Ec*CRP has a very strong preference for T_6_ over C_6_ because T_6_ allows *Ec*CRP to kink DNA as part of forming a strong and specific protein-DNA interaction [23]. Even though *Ec*CRP can bind weakly to CRP-S sites *in vitro* without the co-activator Sxy, *Ec*CRP binding alone is insufficient for transcriptional activation of gene promoters with CRP-S sites [22]. Altogether, gene expression and protein-DNA interaction experiments led us to hypothesize that Sxy enhances CRP binding at CRP-S sites, perhaps by stabilizing CRP dimer formation. As well, Sxy may assist in recruitment of RNA polymerase to gene promoters where CRP is weakly associated with CRP-S sites.

Sxy’s role as the central activator of natural competence was discovered in *H. influenzae* over 25 years ago [4]. Sxy’s role as an activator of CRP-S promoters is conserved across three Gamma-proteobacteria families: Enterobacteriaceae, Pasteurellaceae, and Vibrionaceae, [1,5,7,9,10,18,22,24–27], yet Sxy’s mode of action remains unknown. Unfortunately, *in vitro* characterization of Sxy has been hampered by the toxicity of overexpressed *sxy* and the requirement for strong denaturants to solubilise Sxy inclusion bodies [7,15,25]. We originally proposed that Sxy’s role might be to bind to A+T sequences upstream of *H. influenzae* CRP-S sites [22], but similar sequences were not detected upstream of *E. coli* CRP-S sites [7]. Previous studies from our laboratories indicated that CRP and Sxy have conserved functions in *E. coli* and *H. influenzae*. First, we found that *Ec*CRP can activate competence gene expression in a *H. influenzae* Δ*crp* mutant, and that this complementation absolutely requires Sxy [9, 28]. Second, reciprocal experiments demonstrated that CRP-S promoter activity is higher when cognate pairs of *Ec*CRP/*Ec*Sxy and *Hi*CRP/*Hi*Sxy were co-expressed compared to creating pairs between proteins from different species [7]. These findings suggest that CRP and Sxy work best with their co-evolved protein partner, supporting a model in which these proteins physically interact *in vivo*.

Bioinformatic homology searches (HHSearch and SCOOP) at the Pfam proteins family database predict two putative domains in Sxy: TfoX-N (PF04993) and TfoX-C (PF04994) [29] (Fig. 1A). An unpublished crystal structure is available for TfoX-N from the *sxy* homolog VP1028 of *Vibrio parahaemolyticus* [Protein Data Bank: 2od0], which is predicted to form a dimer. The CATH database of Protein Data Bank hierarchies clusters the TfoX-N domain with several methyltransferases and an acetyltransferase in Superfamily 3.30.1460.30 - YgaC/TfoX-N like chaperone [30]. TfoX-C is predicted to belong to a superfamily of DNA-binding helix-hairpin-helix domains that includes phage lambda’s DNA polymerase and the C-terminus of bacterial RNA polymerase alpha subunit. These bioinformatic predictions are based on hidden Markov models capable of detecting homology between distantly related proteins, thus provide useful material for hypothesis generation but do not provide direct evidence of protein function. Altogether, bioinformatic predictions suggest that the Sxy N-terminal domain could participate in dimerization while the C-terminus is a DNA binding domain; this domain architecture is similar to CRP, where CRP’s effector-binding N-terminal domain is responsible for dimerization while the C-terminal domain is responsible for DNA binding. Also similar are the sizes of *Ec*Sxy (209 amino acids) and *Ec*CRP (210 amino acids).

**Fig. 1.**
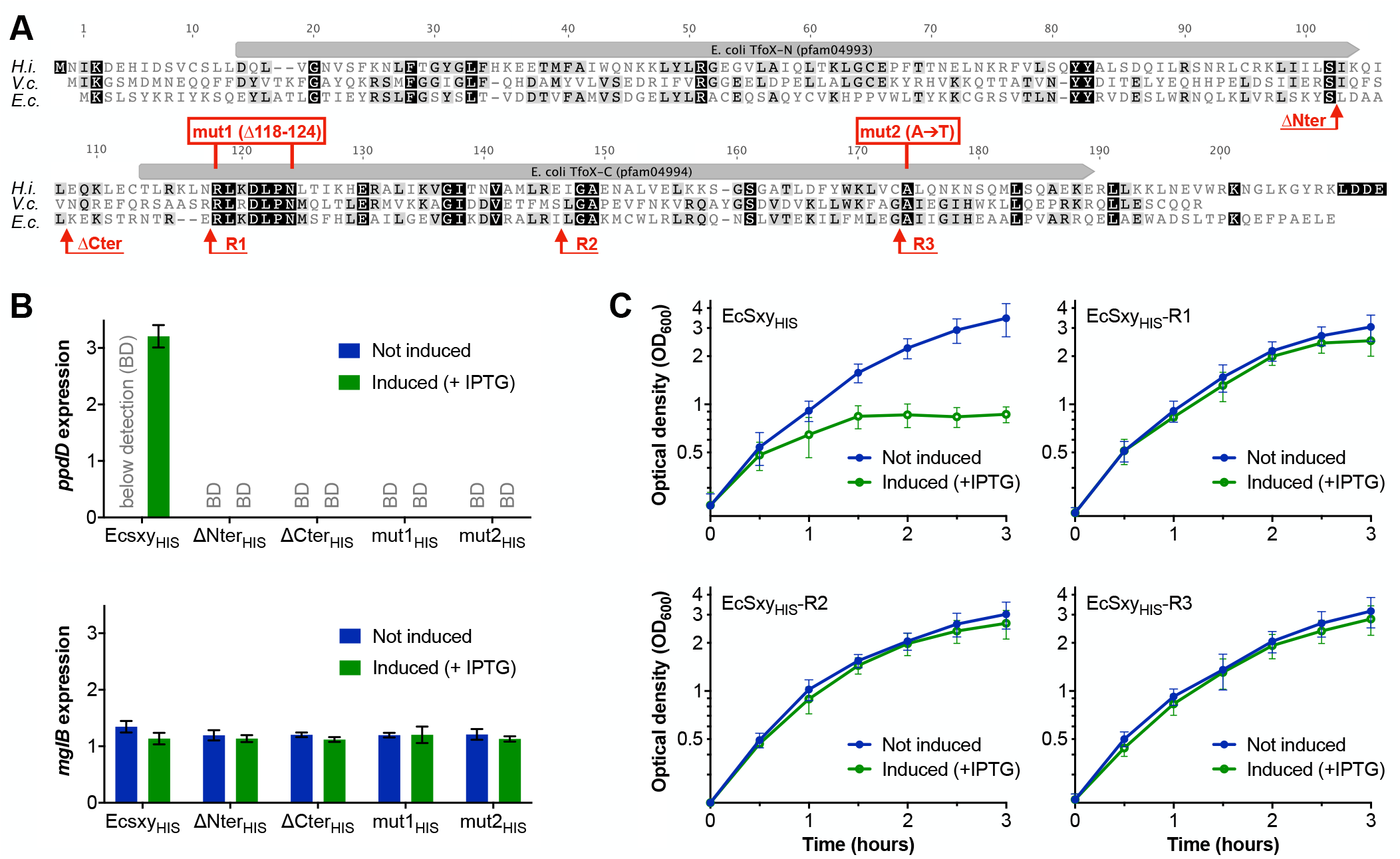
Protein regions and amino acids required for *E. coli* Sxy function. (A) Alignment of Sxy protein sequences from *E. coli* (b0959), *H. influenzae* (HI0601) and *V. cholerae* (VC1153). Amino acids that are identical across all aligned regions are highlighted black, while similar amino acids are highlighted grey. Position numbering is according to *Ec*Sxy. Alignment of all mutant sequences is provided as S1 Fig. (B) Quantitative PCR (qPCR) measurement of changes in the expression of *pilA* (*ppdD*) and *mglB* 60 minutes after IPTG induction of plasmid-encoded *sxy* gene variants. Mean and range of two biological replicates are plotted. Expression levels were normalised by diluting RNA samples 1:10,000 and quantifying 23S rRNA levels; thus the y-axis indicates expression levels approximately 1:10,000 of 23S rRNA expression levels. BD, below detection. (C) The growth inhibition phenotype characteristic of *Ec*Sxy_HIS_ overexpression is reduced by removing amino acids from the Sxy C-terminus. Mean and standard deviation of five biological replicates are plotted.

Here we report multiple experimental approaches designed to characterize how Sxy might participate in CRP-Sxy and Sxy-DNA interactions. Phenotypic and gene expression assays were useful for confirming the requirement for specific regions of the Sxy protein. However, assays relying on protein purification yielded promising but ultimately misleading evidence of protein-protein and protein-DNA interactions. Thus, we present a cautionary report and detail the need for extensive confirmatory experiments when testing for putative Sxy-CRP and Sxy-DNA interactions.

## Results and Discussion

### Diverse mutations in Sxy prevent CRP-S promoter activity and relieve growth inhibition

To identify amino acids and domains required for Sxy’s conserved function as an activator of competence gene expression in the Enterobacteriaceae, Pasteurellaceae, and Vibrionaceaea, we aligned the Sxy protein sequence from a representative member of each genus. This identified conserved amino acids potentially important for Sxy function (Fig. 1A). All three Sxy proteins have similar lengths (209, 215, and 199 amino acids, respectively), but amino acid sequence identity (21-26 %) was low and distributed throughout their lengths. Each was also predicted to contain both the TfoX-N and TfoX-C domains (domains are illustrated in Fig. 1A). To test whether either predicted domain is sufficient for transcriptional activation of a CRP-S regulated gene, we deleted the N-terminal half of *Ec*Sxy to amino acid 102 (*Ec*Sxy_HIS_-ΔNter) and the C-terminal half from position 108 (*Ec*Sxy_HIS_-ΔCter) (Fig. 1A). We also tested the requirement for the largest block of conserved amino acids in all three genera by deleting the eight amino acids between positions 118 and 125, creating mutant *Ec*Sxy_HIS_-mut1. Alanine (Ala) at position 174 is conserved in 98% of the 1,306 full-length Sxy orthologs annotated at EMBL InterPro; we converted alanine 174 to threonine, creating *Ec*Sxy_HIS_-mut2 (Fig. 1A).

To assess how each mutation in *Ec*Sxy impacted transcription activation, we measured transcriptional activity at the *pilA* (*ppdD*) CRP-S promoter and the *mglB* CRP-N promoter. Expression of *pilA* was only detected when wildtype *Ec*Sxy_HIS_ was over-expressed, whereas all Sxy mutant variants were incapable of inducing *pilA* (*ppdD*) expression (Fig. 1 B). Conversely, expression of *mglB* was unaffected by overexpression of wildtype or Sxy variants (Fig. 1 B). Altogether, the TfoX-N and TfoX-C domains, amino acids 118-125, and Ala174 were all critical for transcriptional activation of *pilA*. These various mutations in Sxy may prevent *pilA* expression either because transcription activation requires Sxy-CRP interactions that are lost in the mutant proteins, and/or because the mutations abolish an independent function for Sxy at the *pilA* promoter.

Another phenotype associated with Sxy arises when *sxy* is overexpressed in *E. coli*. Overexpression of cloned *sxy* causes growth inhibition and induction of the RpoH stress response, and this growth inhibition phenotype manifests even at low concentrations of the inducer, IPTG [5, 7]. We systematically truncated the C-terminus of histidine-tagged Sxy (*Ec*Sxy_HIS_) and tested for the growth inhibition phenotype. Sxy inhibition of growth was alleviated by all three C-terminus truncates, *Ec*Sxy_HIS_-R1, *Ec*Sxy_HIS_-R2, and *Ec*Sxy_HIS_-R3 (truncates length indicated in Fig. 1A, and growth is illustrated in Fig. 1C). IPTG-induced expression of the truncated proteins was confirmed by mass spectrometry (described below), indicating that the full-length C-terminus is required for growth inhibition. When not induced by IPTG, cells were unaffected by cloned wildtype and mutant *sxy* genes (S1 Fig. B).

### Sxy does not enhance CRP activation of CRP-N-regulated carbon metabolism genes

Next, we examined whether Sxy impacts CRP-regulated phenotypes other than natural competence and growth inhibition. Specifically, because *Ec*CRP has a higher affinity for DNA than *Hi*CRP does [22], we used the CRP-regulated phenotypes of carbon metabolism to test whether Sxy enhances CRP-induced gene expression at genes with CRP-N sites. CRP activates the uptake and metabolism of diverse carbon sources by *E. coli* and *H. influenzae*. In *E. coli*, CRP was required for metabolism of maltose, mannitol, xylose and glycerol, but not fructose or galactose (S1 Table, rows 1 and 2), and exogenous expression of *Ec*CRP_HIS_ from plasmid p*Eccrp* fully restored wildtype metabolism to a Δ*crp* mutant (S1 Table 1, row 3). In contrast, complementation by *Hi*CRP_HIS_ was partial, restoring only xylose metabolism (S1 Table 1, row 4). Suggesting that *Hi*CRP_HIS_’s lower affinity for DNA prevents it from activating all *E. coli* CRP-N promoters. We tested this by examining whether *Hi*Sxy_HIS_ could restore maltose, glycerol, and mannitol fermentation by stabilizing *Hi*CRP_HIS_-DNA interactions. Although co-expression of *Hi*CRP_HIS_ and *Hi*Sxy_HIS_ restores CRP-S promoter function in *E. coli* [7], the same co-expression did not restore fermentation of maltose, glycerol, and mannitol in our phenotypic assays (S1 Table 1, rows 5 and 6). Further, *Ec*CRP activity at CRP-N promoters was unaffected by expression of either *Ec*Sxy_HIS_ or *Hi*Sxy_HIS_ (S1 Table 1, rows 7 and 8), suggesting that Sxy cannot significantly enhance the binding of CRP to weak CRP-N sites sufficiently to enhance transcription.

### Protein-DNA crosslinking *in vivo* detects non-specific DNA binding

We next sought to test whether Sxy-DNA interactions occur *in vivo* using a chromatin affinity precipitation assay. In principle, formaldehyde can crosslink *Ec*Sxy_HIS_ with DNA and proteins to which it is bound *in vivo*. Isolation of *Ec*Sxy_HIS_ on a nickel affinity column will then co-purify any DNA bound and crosslinked to *Ec*Sxy_HIS_. To test whether Sxy physically interacts with DNA in vivo, the cloned *Ecsxy*_HIS_ gene was overexpressed in *E. coli*, formaldehyde was added to cell cultures to crosslink interacting DNA and protein molecules, and cells were lysed by sonication and fractionated by centrifugation. The cytoplasmic (soluble) fraction was then incubated with nickel-agarose resin (Ni-NTA) to bind *Ec*Sxy_HIS_ and any crosslinked DNA. After extensive washing, *Ec*Sxy_HIS_ was eluted with imidazole, formaldehyde crosslinking was reversed, and eluted DNA was quantified by PCR.

We hypothesized that Sxy bound specifically to CRP-S sites in living cells, which we could detect as an enrichment of CRP-S containing loci compared to non-CRP-S containing DNA in elution fractions. We quantified the levels of three unlinked chromosomal genes: two CRP-S regulated genes, *pilA* (*ppdD*) and *comM*, and a negative control non-CRP regulated gene *hns*. In each fraction, all three genes were eluted in equal quantities (Fig. 2, upper panel), indicating that DNA retention was not specific to genes with CRP-S sites. In the absence of formaldehyde crosslinking *in vivo*, most DNA was eluted in fraction #1 and DNA concentrations generally declined in subsequent fractions (Fig. 2, lower panel). This elution pattern did not correspond to protein elution, confirming that formaldehyde crosslinking contributed to DNA retention on nickel affinity columns.

**Fig. 2.**
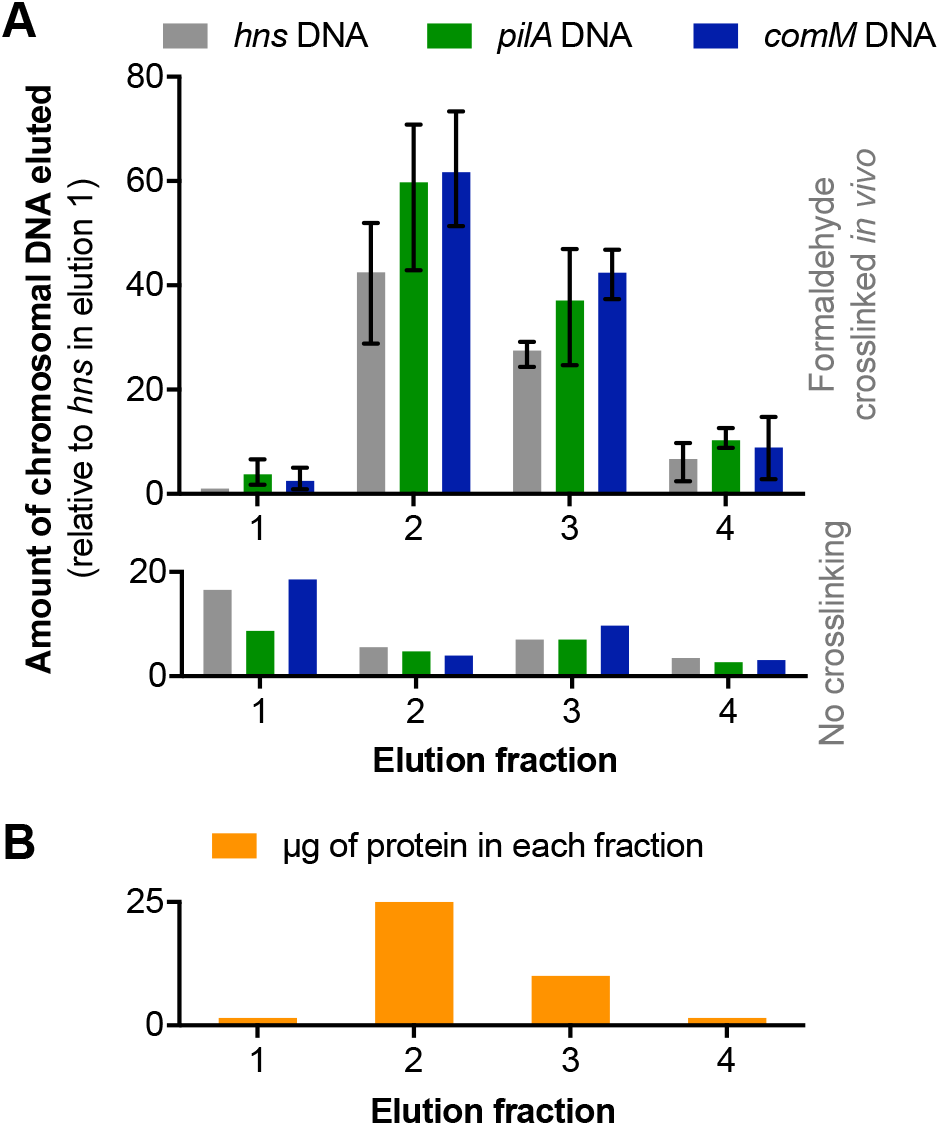
Co-elution of DNA with protein fractions containing *Ec*Sxy_HIS_. (A) qPCR measurement (in arbitrary units) of DNA eluted from nickel affinity columns. The mean and range of two biological replicates are plotted for formaldehyde crosslinked samples (upper panel) and a single biological sample without crosslinking (lower panel). (B) Total µg of protein eluted in each fraction.

### Quantifying CRP-DNA and Sxy-DNA interactions *in vitro*

We have long hypothesized that Sxy is required for transcriptional activation at CRP-S promoters because it stabilizes CRP-DNA interactions [22]. Electrophoretic mobility shift assays (Bandshift assay), in which protein-DNA binding retards electrophoretic migration of bait DNA [22], are useful for detecting protein-DNA interactions and for quantifying binding affinity. *Ec*CRP_HIS_ and *Hi*Sxy_HIS_ were purified in their native forms as in earlier studies [22]. Expression of *HiSxy*_HIS_ in *E. coli* for purification of *Hi*Sxy_HIS_ causes formation of inclusion bodies [25]. Nevertheless, expression of *HiSxy*_HIS_ and *EcSxy*_HIS_ produces sufficient soluble protein to generate the phenotypes observed in Figures 1 and 2, and in past publications [5, 7]. Thus, nickel affinity columns were used to purify this native (soluble) *Hi*Sxy_HIS_ and *Ec*Sxy_HIS_, and coomassie-stained polyacrylamide gel electrophoresis and western blotting confirming the isolation of Sxy_HIS_ proteins [25].

To test for Sxy-DNA and Sxy-CRP-DNA interactions, we designed protein-DNA binding reactions to distinguish between CRP’s lower affinity for DNA fragments containing a native CRP-S promoter (*H. influenzae pilA*) and its higher affinity for a derivative CRP-N promoter created from it by site-directed mutagenesis (*pilA*-N) (Fig. 3 A) [22]. As previously demonstrated, *Ec*CRP_HIS_ bound both *pilA* and *pilA*-N sites *in vitro*, shifting 30% of the low affinity *pilA* and 77% of *pilA*-N with 2 nM *Ec*CRP_HIS_ (Fig. 3 B and C, lane 2). Also as previously shown [22], *Hi*CRP_HIS_ did not bind the native *pilA* site, and required 20 nM of *Hi*CRP_HIS_ to shift only 6% of *pilA*-N DNA (Fig. 3 E and F, lane 2).

**Fig. 3.**
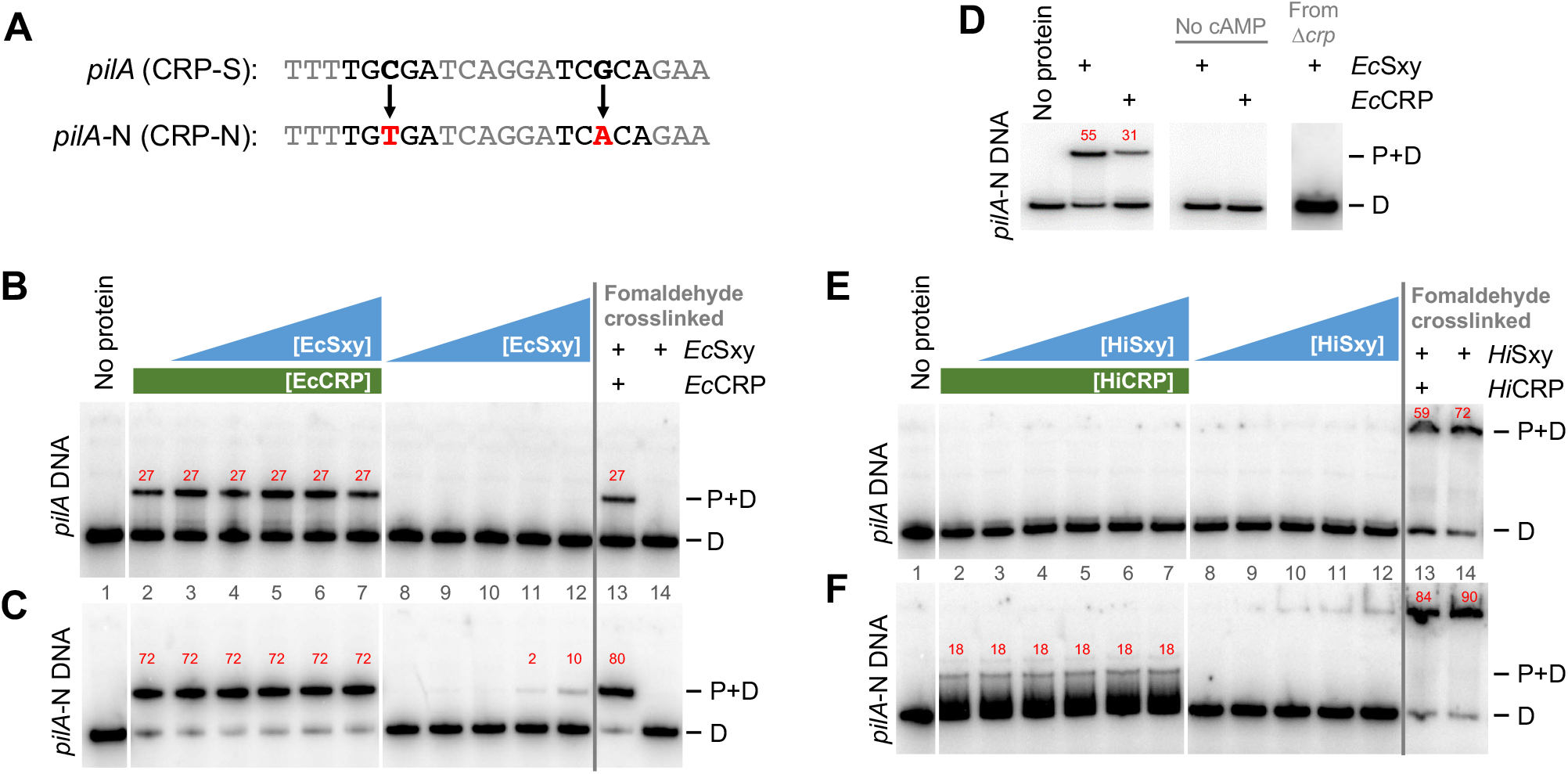
Binding of *E. coli* and *H. influenzae* CRP and Sxy proteins to promoter DNA. (A) CRP binding site sequences in bait DNA. Arrows and bold lettering indicate the point mutations that converted the native *H. influenzae pilA* sequence to *pilA*-N. (B, C, E, F) Bandshift analysis of CRP (green) and Sxy (blue) binding to *pilA* or *pilA*-N DNA. Red numbers indicate the percentage of shifted DNA. Each panel is a single gel, with white and grey lines overlaid to facilitate lane identification; “P+D” indicates protein-DNA complexes, “D” indicates DNA alone. In all gels, Lane 1: DNA alone, Lane 2: CRP + DNA, Lanes 3 to 7 and 13: Sxy and CRP mixed together before incubation with DNA, Lanes 8 to 12 and 14: Sxy + DNA. Lanes 13 and 14: samples were crosslinked with formaldehyde before gel loading. (B and C) *E. coli* (*Ec*) proteins. Protein concentrations: 2 nM *Ec*CRP_HIS_ in lanes 2-7 and 13; 0.2, 1, 2, 4, and 20 nM *Ec*Sxy_HIS_ in lanes 3-7 and 8-12, respectively, and 20 nM *Ec*Sxy_HIS_ in lane 14. (D) Bandshift analysis of *Ec*Sxy_HIS_ binding to *pilA*-N DNA with or without cAMP. Lanes 2, 4 and 6: *Ec*Sxy_HIS_ (700 nM) + DNA, Lanes 3, 5 and 7: *Ec*CRP_HIS_ (4 nM) + DNA, Lane 3: DNA alone. Lanes 4 and 5: no cAMP, Lanes 1 to 3 and 6: with cAMP. Lanes 2 and 4: *Ec*Sxy_HIS_ purified from wildtype cells, Lane 6: *Ec*Sxy_HIS_ purified from Δ*crp* cells. The uncropped gel is provided in Supporting Material S2 Figure. (E and F) *H. influenzae* (*Hi*) proteins. Protein concentrations: 20 nM *Hi*CRP_HIS_ in lanes 2-7 and 13; 2, 10, 20, 40, and 200 nM *Hi*Sxy_HIS_ concentration in lanes 3-7 and 8-12, respectively, and 20 nM *Hi*Sxy_HIS_ in lane 14.

The different affinities of *Ec*CRP_HIS_ and *Hi*CRP_HIS_ for DNA allowed us to test whether *E. coli* or *H. influenzae* Sxy could enhance binding of their cognate CRP to *pilA* or *pilA*-N DNA. Enhanced binding was predicted to manifest as an increase in the amount of *pilA* DNA shifted by *Ec*CRP, and to create a detectable shift of *pilA* DNA when bound by *Hi*CRP. Additionally, we predicted that simultaneous binding of CRP and Sxy to DNA would create a super-shift; in other words, a CRP-Sxy-DNA complex is expected to migrate slower during electrophoresis than the smaller CRP-DNA complex. However, a range of Sxy concentrations had no detectable effect on either *Ec*CRP_HIS_ or *Hi*CRP_HIS_ binding to *pilA* or *pilA*-N DNA (Fig. 3 B, C, E, F, lanes 3-7).

We next tested whether *Ec*Sxy_HIS_ or *Hi*Sxy_HIS_ alone can bind DNA. Surprisingly, at 4 and 40 nM of *Ec*Sxy_HIS_, a small amount of *pilA*-N DNA was shifted, producing a new band at the same position as CRP-DNA binding (Fig. 3 C, lanes 11 and 12). The shift was dependent on CRP’s allosteric effector cAMP, confirming the presence of CRP in the binding reactions (Fig. 3 E, compare lanes 2 and 4). No binding was detected when the experiment was repeated using *Ec*Sxy_HIS_ purified from a Δ*crp* background, confirming that the shift arose from contaminating CRP in the *Ex*Sxy_HIS_ preparation (Fig. 3 D, lane 6). Comparison with lane 11 indicates that CRP contamination at <100 pM concentration would be sufficient to account for the observed shift (Fig. 3 C, lanes 2 and 12). With *Hi*Sxy_HIS_ and *pilA*-N bait DNA, increasing protein concentrations correlated with an increase in the amount of DNA retained in the gel wells (Fig. 3 F, lanes 8-12). A specific interaction between Sxy and a DNA binding site was expected to yield approximately the same sized bandshift as CRP-DNA binding, but no such band was detected, suggesting *Hi*Sxy_HIS_ was interacting non-specifically with *pilA*-N DNA to retain it in the gel wells.

To test whether Sxy-DNA binding may in fact occur but is too weak to persist under electrophoretic conditions, we used formaldehyde to cross-link proteins bound to DNA *in vitro*, before electrophoresis. A 1:1 mix of *Ec*CRP_HIS_:*Ec*Sxy_HIS_ resulted in a shift corresponding in size to a CRP-DNA complex (Fig. 3 B and C, lanes 13). In contrast, with the *H. influenzae* proteins, most bait DNA was retained in the wells of the gels, indicating the formation of large protein-DNA complexes (Fig. 3 E and F, lanes 13 and 14). This super-shift occurred also in lane 14, which contained only Sxy without CRP, supporting the hypothesis that DNA-Sxy interactions exist, and the super-shift is evidence that Sxy-DNA interactions could be stabilized with formaldehyde.

We also tested the hypothesis that CRP and Sxy may need to form protein-protein contacts prior to binding DNA. However, pre-mixing the proteins from each species before adding bait DNA had no effect on DNA binding (S3 Fig., lanes 4 and 10). Similarly, combining either CRP or Sxy alone with bait DNA before addition of the second protein also had no effect (S3 Fig., lanes 5, 6, 11 and 12). Collectively, these results confirm that Sxy does not bind stably to DNA *in vitro*, nor does it enhance CRP binding to DNA in bandshift assays. Nevertheless, the presence of contaminating CRP in *Ec*Sxy_HIS_ preparations was suggestive of protein-protein interactions between Sxy and CRP.

Lastly we tested if very high concentrations of *Ec*Sxy_HIS_ isolated from Δ*crp* cells could shift *pilA* DNA (the highest Sxy concentration used in the above-mentioned experiment (Fig. 3) was 2nM). Purified *Ec*Sxy_HIS_ proteins (10 mM) shifted 17 % of DNA (Fig. 4 A, lane 3), and higher *Ec*Sxy_HIS_ concentrations resulted in stronger DNA shifts (40%, 64%, and 74 %) (Fig. 4 A, lane 4 - 6). The discrete bandshift supported that this was specific protein-DNA interactions. Further, an excess of non-specific competitor DNA (poly dI-dC) only caused a minor reduction in the observed protein-DNA interaction (Fig. 4 A, lane 2). Titrating salmon sperm DNA as a non-specific competitor revealed that once competitor DNA surpassed the concentration of bait DNA (around 7 ng/reaction), protein binding to bait DNA was reduced (Fig. 4 B). This suggested that although a discrete bandshift was formed by adding *Ec*Sxy_HIS_ extract to bait DNA, protein-DNA binding was low affinity and/or low specificity. Adding high concentrations of *Ec*CRP_HIS_ to *Ec*Sxy_HIS_ reactions produced an additive effect resulting in protein-DNA supershifts (S6 Fig., lanes 3, 4, 8, and 9). The retention of bait DNA in wells is consistent with CRP proteins binding non-specifically to bait DNA. Non-specific protein-DNA interactions were confirmed as competitor DNA effectively liberated DNA from wells (S6 Fig., lanes 5 and 6).

**Fig. 4.**
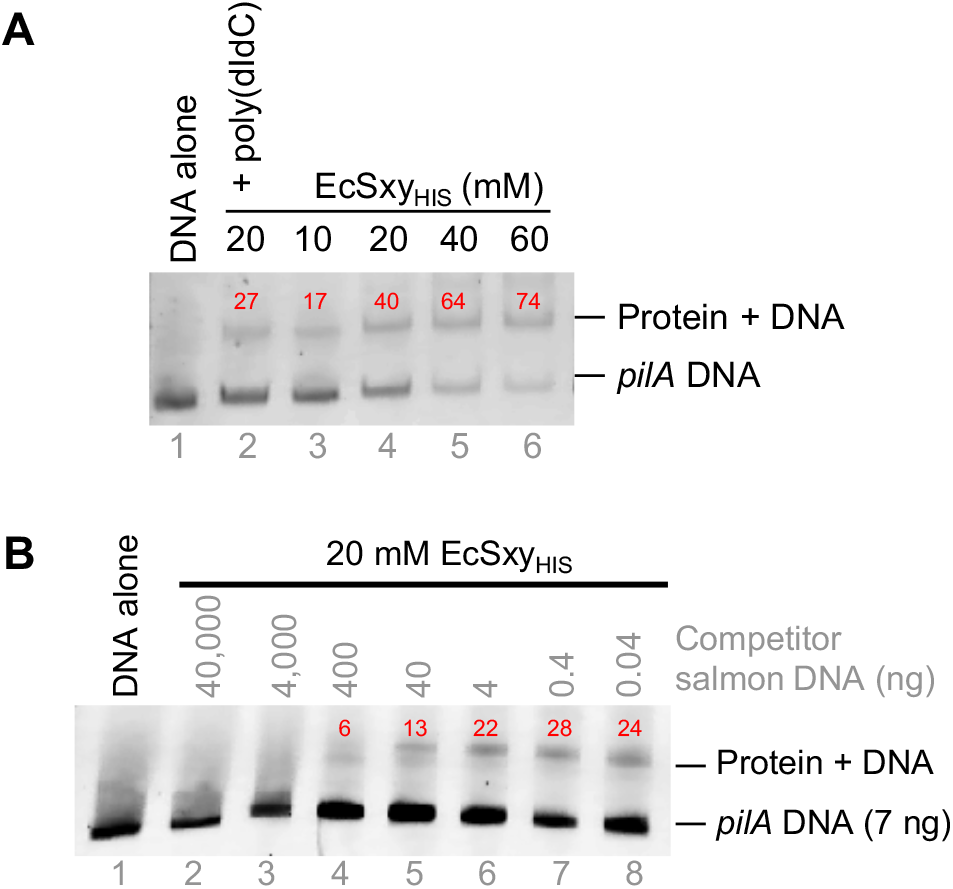
High concentration *Ec*Sxy_HIS_ protein extract in bandshift assays. (A) Titration of *Ec*Sxy_HIS_ extract from Δ*crp* cells. (B) Titration of non-specific competitor salmon sperm DNA. *Ec*Sxy_HIS_ was purified from Δ*crp* cells and incubated with *pilA* DNA. Red numbers indicate the percentage of shifted DNA. Uncropped gels are provided in Supporting Material S4 Figure and S5 Figure.

The results above indicate that *Ec*Sxy_HIS_ and *Hi*Sxy_HIS_ extracts contain non-specific DNA binding activity that can be detected by bandshift assays either by stabilizing protein-DNA interactions with formaldehyde (Fig. 3) or by addition of high concentrations of protein extract (Fig. 4 and S6 Fig.).

### Identification of other DNA binding proteins in Sxy extracts

Previous studies of protein purification by immobilized metal affinity chromatography identified SlyD and CRP from native *E. coli* extracts to have a high affinity for nickel affinity columns, and that these contaminating proteins elute at high imidazole concentrations along with the desired histidine-tagged protein [31, 32]. The SlyD protein chaperone contains a metal binding domain with high affinity for Zn^2+^ and Ni^2+^ ions, making it an expected contaminant of metal affinity chromatography. We used mass spectrometry to examine the degree to which *Ec*Sxy_HIS_ was enriched by nickel affinity column purification, also allowed us to identify and quantify cytoplasmic proteins co-purifying with *Ec*Sxy_HIS_.

We first established that *Ec*Sxy_HIS_ was not detected in the cytoplasm of *E. coli* Δ*crp* cells before IPTG induction, then became the 4^th^ most abundant protein in cytoplasm after induction (S7 Fig.). After nickel column purification, *Ec*Sxy_HIS_ represented almost half of the eluted protein (40%), but was surpassed by SlyD (43%) (Fig. 5A). In the eluate from wildtype cultures, *Ec*Sxy_HIS_ was only the 4^th^ most abundant protein (6% of eluted protein) (Fig. 5B). *Ec*Sxy_HIS_-R3 only represented 3% of eluted protein when enriched from wildtype cells (Fig. 5C). The DNA-binding proteins Fur and CRP were the 2^nd^ and 3^rd^ most abundant proteins in eluates from wildtype cells (Fig. 5B, C, D). The presence of Fur, CRP and other DNA-binding proteins in all fractions suggests that the observations of putative Sxy-DNA and Sxy-CRP interactions by bandshift assays were in fact artefactual.

**Fig. 5.**
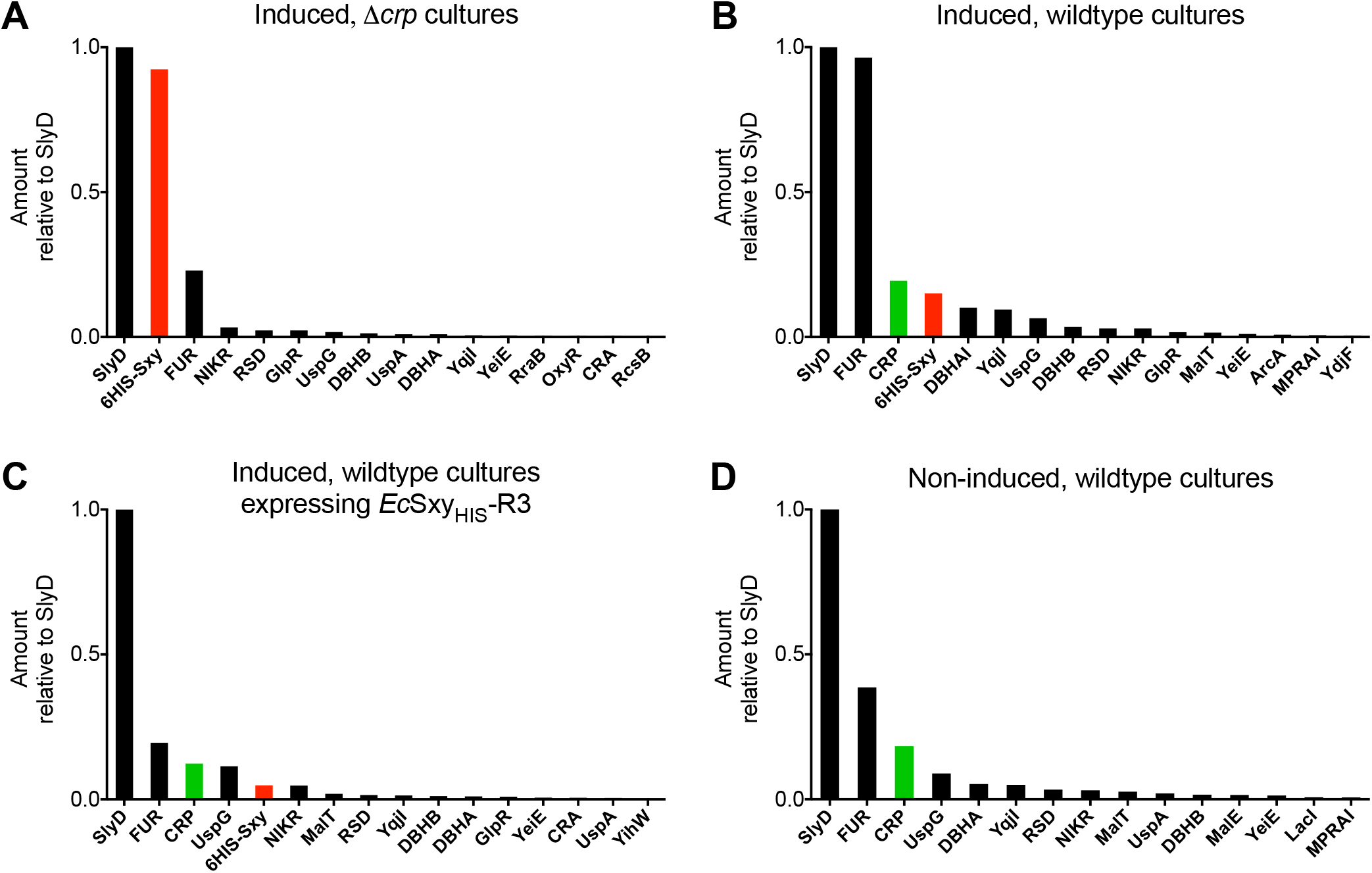
Mass spectrometry detection of *Ec*Sxy_HIS_ in Ni-NTA column eluates. (A) Eluate from *E. coli* Δ*crp* carrying p*Ecsxy* and induced with IPTG. (B) Eluate from wildtype *E. coli* carrying p*Ecsxy* and induced with IPTG. (C) Eluate from wildtype *E. coli* carrying p*Ecsxy*-R3, which produces *Ec*Sxy_HIS_-R3. (D) Eluate from wildtype *E. coli* carrying p*Ecsxy*. The eluates in A & B correspond to *Ec*Sxy_HIS_ samples used in bandshifts. In all graphs, the 16 most abundant proteins are ranked from high to low abundance; *Ec*Sxy_HIS_ is highlighted in red and native CRP is highlighted in green.

## Conclusions

The present work used protein alignments, directed mutations, and phenotypic assays to confirm that Sxy’s predicted domains and conserved amino acids are essential for transcriptional activation of a CRP-S promoter in *E. coli*. Growth assays confirmed that growth inhibition by Sxy requires a full-length C-terminus, which is predicted to encode a helix-hairpin-helix DNA binding domain. All experiments described in this work (both *in vivo* and *in vitro*) used histidine-tagged Sxy proteins, and multiple lines of evidence suggest that the histidine tag does not negatively impact Sxy function. For example, overexpression of either wildtype or histidine-tagged Sxy inhibits *E. coli* growth, and both histidine-tagged Sxy and CRP are strong activators of gene expression from CRP-S promoters (Fig. 1B and in [7, 22]). Having confirmed these phenotypes resulting from overexpressing histidine-tagged *sxy* genes, we tested the hypothesis that Sxy enhances weak CRP-DNA interactions using *Hi*CRP’s inability to fully restore sugar metabolism in an *E. coli* Δ*crp* mutant (S1 Table). However, the observed negative results are inconclusive because they may represent an insurmountable inability of *Hi*CRP to act at all *E. coli* CRP-N promoters. Nevertheless, the negative results in the sugar metabolism assys could also reflect that CRP-N promoters lack a specific, but as yet undetected, feature required for Sxy activity.

Nickel affinity pull-down assays and bandshift assays appeared to demonstrate DNA binding by *Ec*Sxy_HIS_, but the DNA binding activities are best explained by the activity of contaminating proteins in nickel affinity column purification. Thus, the complication of bandshift and pull down assays by contaminating DNA-binding activity provides a valuable warning of the need for extensive controls and confirmatory experiments until robust *in vitro* assays for Sxy activity are identified. Throughout this study we applied all standard quality control steps and performed preparative techniques as used successfully for other bacterial proteins. Hence, we present this study partly as a cautionary tale against the use of nickel affinity purification to isolate a low-concentration protein. Even though Ni-NTA resin columns are widely used to purify histidine-tagged proteins [33–35], we discovered many metal binding and other co-purified proteins eluted in this system despite extensive washing before elution. Therefore, we recommend using mass spectrometry or other precision techniques to assess the purity of proteins eluted from Ni-NTA systems.

Despite the artefactual and ultimately negative results for protein-protein-DNA interactions presented above, the simplest model that remains consistent with all genetic and phenotypic data is that Sxy-CRP interactions facilitate CRP binding to DNA and transcriptional activation at CRP-S promoters. To bind DNA and recruit RNA polymerase, CRP must dimerize and then deform DNA by kinking [21]. Although CRP-S sites are targets for CRP binding [22], their characteristic sequence reduces CRP’s ability to bend and kink DNA [23]. There may be a role for Sxy in strengthening CRP dimerization, thus stabilizing CRP to spend sufficient time in association with CRP-S sites to achieve the required DNA bending and RNA polymerase recruitment. The CRP-S regulon is unique among characterized bacterial promoter mechanisms because it is defined by a distinct DNA element (the CRP-S site) embedded within the binding site of a global transcription factor, CRP. The molecular interactions that connect Sxy, CRP, and CRP-S sites to stimulate transcription remain enigmatic; additional phenotypes and alternate purification strategies are required to identify Sxy’s cellular function(s).

## Materials and Methods

### Bacterial strains, plasmids, and growth conditions

*E. coli* BW25113 and JW5702 (*crp::*kan) were obtained from the KEIO collection [36]. For protein purification, *sxy*_HIS_ and *crp*_HIS_ were expressed in *E. coli* BL-21 from p*Ecsxy*, p*Hisxy*, p*Eccrp* and p*Hicrp*, which are detailed in [9] and [22]. Gene expression, growth, and carbon metabolism assays were conducted in *E. coli* W3110.

*E. coli* was grown on Luria Bertani (LB) broth or LB agar (1.2 or 1.5 %) at 37 °C. When required, antibiotics were used at the following concentrations: kanamycin 15 μg/ml, chloramphenicol 20 μg/ml, ampicillin 100 μg/ml, tetracycline 10 μg/ml. 1 mM of Isopropyl β-D-1-thiogalactopyranoside (IPTG) was used to induce plasmid-encoded gene expression. Complementation of the Δ*crp* carbon metabolism phenotypes was assessed by streaking on Difco MacConkey indicator plates containing 1 % of the test sugar; fermentation of the test sugar resulted in pink colonies.

### Creation of *sxy* mutations and truncations

Directed mutations and truncations in *E. coli sxy* were created by inverse PCR using p*Ecsxy* as a template, then were confirmed by restriction digest and Sanger DNA sequencing. All mutations maintain the *sxy* coding sequence in frame and retain the N-terminal hexa-histidine tag (primers are listed in S2 Table).

### Quantitative PCR

Quantitative PCR was performed as described in [1], where RNA was extracted using RNeasy Mini Kits (Qiagen), followed by DNase treatment with a DNA Free kit (Ambion). cDNA templates were synthesized using the iScript cDNA synthesis kit (BioRad). qPCR was carried out using SYBR Green Supermix (BioRad) with the primers listed in S2 Table.

### Sxy growth inhibition assays

*E. coli* W3110 containing either plasmid p*Ecsxy* (WT), p*Ecsxy*-R1, p*Ecsxy*-R2, or p*Ecsxy*-R3 was cultured in 40 ml LB with 35 μg/ml chloramphenicol in a shaking incubator at 37 °C. At OD_600_ of 0.2, the culture was separated into two flasks: one induced with 1mM IPTG and one control culture “non-induced”. Growth was monitored for 4 hours after IPTG induction by measuring light absorbance (OD_600nm_).

### Protein purification

The histidine-tagged CRP proteins (25 KDa) are described in Cameron and Redfield [22]. The histidine-tagged Sxy proteins (26 KDa) were purified under the same conditions after 2.5 hours expression time after 1 mM IPTG induction at OD_600_ 0.5. Cells were harvested (3,000 × g, 4 °C, 20 minutes) and cell pellets were frozen at - 20 °C overnight. Frozen cell pellets were thawed on ice, re-suspended in 2 ml protein lysis buffer (50 mM NaH_2_PO_4_, 200 mM NaCl, 10 mM imidazole, pH 8.0) to which 1 X of protease inhibitor (Pierce™ Protease Inhibitor Tablets, EDTA-free from Thermo Fisher) was added and then incubated at 4 °C for 1 hour with rotation. Cell suspensions were further lysed by sonication in 15 ml falcon tubes with a Bioruptor® Standard (Diagenode) (Power high, 30 sec ON/30 sec OFF for 15 minutes at 4 °C). Cell debris was removed by centrifugation (3,000 × g, 4 °C, for 20 minutes) and the supernatant saved for protein purification. Purification of His-tagged proteins was carried out using Qiagen polypropelene columns (Ni-NTA resin) according to the manufacturer guidelines. After three washes with wash buffer (50 mM NaH_2_PO_4_, 300 mM NaCl, 250 mM imidazole; pH 8), His-tag proteins were eluted from columns by addition of 2 ml elution buffer (50 mM NaH_2_PO_4_, 200 mM NaCl, 250 mM imidazole, pH 8.0) and collected in four elution fractions (E1 = 250 µl, E2 =500 µL, E3= 250 µL, E4 = 1 ml).

Eluted protein was centrifuged in 10 kDa filters (Microcon) for 5 minutes at 9,000 × g at 4 °C. 100 µl of storage buffer (20% glycerol, 40 mM Tris, 200 mM KCl) was added to ∼10 µl of filtered proteins. Protein samples were frozen at −80 °C. Protein concentrations were determined by either Bradford assay, BCA assay, Pierce™ BCA Protein Assay Kit from ThermoFisher (product # 23223), or 660 nm assay using Pierce™ 660nm Protein Assay Reagent from ThermoFisher (product prod # 1861426). To confirm protein expression, protein samples were denatured in Laemli sample buffer then heated for 5 minutes at 100 °C, followed by electrophoresis on 12% SDS-PAGE gels, and stained with Coomassie blue.

### Bandshift experiments

Bait DNAs in Fig. 3 and S3 Fig., and their preparation are described in Cameron and Redfield [22]. Bait DNAs in Fig. 4 and their preparation are described in [38]. To make the fluorophore-labelled baits, the promoters of *pilA*, *ppdA* (also called *comN)*, and *mglB* (positive control) were PCR amplified (268 bp, 239 bp, 275 bp respectively) with M13 sequences added to both 5’ and 3’ ends (primers are listed in S2 Table). Then, PCR products were amplified using Fluorescent Oligonucleotides (Cy5-M13 and FAM-M13). Amplicons were then purified and used as DNA baits in Bandshift experiments.

Protein-DNA reactions (5 μl) contained between 10 and 700 nM protein(s) mixed with 4 nM bait DNA in reaction buffer (8 mM Tris (pH 8.0), 30 mM KCl, 3% (v/v) glycerol, 250 µg/ml bovine serum albumin, 100 µM cAMP, and 1 mM dithiothreitol). Reactions were mixed on ice and then incubated at room temperature for 20 minutes before loading onto a 4 °C running polyacrylamide gel (30:1 acrylamide/bisacrylamide; 0.2× TBE [89 mM Tris, 89 mM borate, and 2 mM ethylenediaminetetraacetic acid (pH 8.3)], 2% glycerol, and 200 µM cAMP; running buffer 0.2× TBE and 100 µM cAMP). After electrophoresis for 2.5 hr at 10 mA, the gel was dried and exposed (45 minutes to overnight) to a phosphor screen. Bands were visualized using either a STORM 860 scanner (GE Healthcare) [22] for T4 Polynucleotide Kinase end labelled DNA, or Typhoon FLA 7000 (GE Healthcare) for Cy5 and FAM end-labelled DNA. Gel analysis and band densitometry calculations were conducted using Image J 1.46r.

### Pull-down assays

50 ml LB broth was inoculated 1:100 from overnight *E. coli* cultures harboring plasmid p*Ecsxy*. At OD_600_ 0.5, expression of *sxy* was induced for 2.5 hours by the addition of IPTG (1 mM final concentration). Cells were harvested (3,000 × g), 4 °C, 20 minutes) and washed in 50 ml of ice-cold PBS. Protein-DNA interactions were chemically cross-linked by the addition of formaldehyde (1 % vol/vol final concentration) to the washed cell suspensions which were then incubated with agitation (100 rpm) for 30 minutes at 4 °C. Further cross-linking was inhibited by the addition of ice-cold 2 M glycine (0.125 M final concentration), cells were harvested (3,000 × g, 4 °C, 20 minutes) and cell pellets were frozen at −20 °C overnight. Frozen cell pellets were thawed on ice, re-suspended in 2 ml protein lysis buffer (50 mM NaH_2_PO_4_, 200 mM NaCl, 10 mM imidazole, pH 8.0) to which lysozyme (1 mg/ml final concentration) and protease inhibitor (1 X) (Pierce) were added and then held on ice-for 1 hour. Cell sonication and protein purification was performed as described above. To reverse formaldehyde cross-linking, 5 M NaCl (0.3 M final conc.) was added to 225 µl of each elution fraction and held at 65 °C for 6 hours. Then, 9 µl Proteinase K (10 mg/ml) was added and fractions were incubated at 45 °C overnight, and 2 µl of salmon sperm DNA (5 mg/ml) was added to each sample just before adding 500 µl Phenol:Chloroform:Isoamyl alcohol (25:24:1). The samples were vortexed and centrifuged at 13,000 × g for 5 minutes at room temperature. The aqueous (top) layer was collected in a fresh 1.5 ml microfuge tubes and 500 µl of chloroform was added to each sample. The samples were vortexed and centrifuged at 13,000 × g for 5 minutes at room temperature. The aqueous layer was transferred to a fresh 2.0 ml microfuge tubes. 5 µg of GlycoBlue (5 mg/ml), 1 µl of salmon sperm DNA (5 mg/ml, Invitrogen) and 50 µl of 3M NaAc (pH 5.2) was added to each sample and mixed well. The DNA was precipitated with 1375 µl of 100% ethanol and incubated at −70 °C for 30 minutes (or −20 °C overnight). The samples were centrifuged at 13000 × g for 20 minutes at 4 °C. The DNA pellets were washed with 500 µl of ice-cold 70% ethanol and air-dried for 10-15 minutes. The DNA pellets were re-suspended in 50 µl of sterile filtered HPLC water. qPCR analysis was performed for each DNA fraction (four elution fractions).

### Protein mass spectrometric analysis

Protein samples were desalted using 10kDa MWCO filters (EMD Millipore) before tryptic digest. Subsequently, the resulting peptides were separated on a Waters Nanoaqcuity nano-LC and analyzed with a Waters Synapt G2 HDMS (Waters Corporation). For the LC, an Acquity UPLC T3HSS column (75 mm x 200 mm) was used and a gradient was run from 3% acetonitrile/0.1% formic acid to 45% acetonitrile in 2 hours. Mass spectrometric acquisition was conducted using data-independent acquisition (MSE) in resolution mode and using leucine-enkephaline as lockspray for mass correction. Resulting spectra were analyzed with the ProteinLynx Global Server version 3.02 (Waters) with a false discovery rate set to 4%.

## Acknowledgments

We would like to express our appreciation of Tyler Boa for excellent technical assistance.

## Author contributions

### Conceptualization

EYA RJR SS ADSC.

### Funding acquisition

RJR ADSC.

### Investigation

EYA SFF T-CC RCC SS ADSC.

### Writing – original draft

EYA RJR SS ADSC.

## Supporting data legends

**S1 Figure.** (A) Alignment of Sxy C-terminus sequences from *H. influenzae* (HI0601), *E. coli* (b0959), and three C-terminus truncate mutants in *Ec*Sxy_HIS_. (B) Growth of *Ec*Sxy_HIS_ mutant clones when not induced.

**S2 Figure.** Uncropped gel from Figure 3D.

**S3 Figure.** Bandshift analysis of *H. influenzae* and *E. coli* CRP and Sxy proteins binding to *pilA*-N DNA with sequential incubation of proteins in binding reactions. Lanes 1 and 11: DNA alone, Lanes 2 and 12: CRP + DNA, Lanes 3 and 10: Sxy + CRP + DNA mixed together at the same time, Lanes 4 and 9: Sxy + CRP mixed together before addition of DNA, Lanes 5 and 8: CRP + DNA mixed together before addition of Sxy, Lanes 6 and 7: Sxy + DNA mixed together before addition of CRP. For *E. coli* proteins (lanes 1 to 6), 350 nM Sxy and 4 nM CRP were used. For *H. influenzae* proteins (lanes 7 to 12), 700 nM Sxy and 400 nM CRP were used.

**S4 Figure.** Uncropped gel from Figure 4A.

**S5 Figure.** Uncropped gel from Figure 4B.

**S6 Figure.** Bandshift analysis for *Ec*Sxy_HIS_ and *Ec*CRP_HIS_ proteins binding to *H. influenzae* promoter DNA. Protein concentrations are indicated in the Figure. *Ec*SxyHIS was purified from Δ*crp* cells. 120 ng of unlabelled competitor salmon DNA or poly (dI-dC) DNA was added to protein-DNA binding reactions where indicated. Red numbers indicate the percentage of shifted DNA.

**S7 Figure. Induction and mass spectrometry detection of *Ec*Sxy_HIS_ in whole-cell lysates of *E. coli* Δ*crp*.** Protein abundance is ranked from high to low, with only the most abundant DNA-binding proteins presented in the graphs. Sxy is highlighted in red.

**S1 Table. Carbon substrate fermentation by *E. coli* Δ*crp* cells complemented with cognate or non-cognate CRP and Sxy proteins.** Complementation was conducted with plasmids p*Eccrp*, p*Hicrp*, p*Hisxy*, or p*Ecsxy*. Fermentation was assessed on Difco MacConkey indicator plates containing 1 % of the indicated carbon source.

**S2 Table. List of primer sequences.**

