## Supplementary figures and images for "Finding reliable phenotypes and detecting artefacts among *in vivo* and *in vitro* assays to characterize the refractory transcriptional activator Sxy (TfoX) in *Escherichia coli*"

### Supplementary Materials

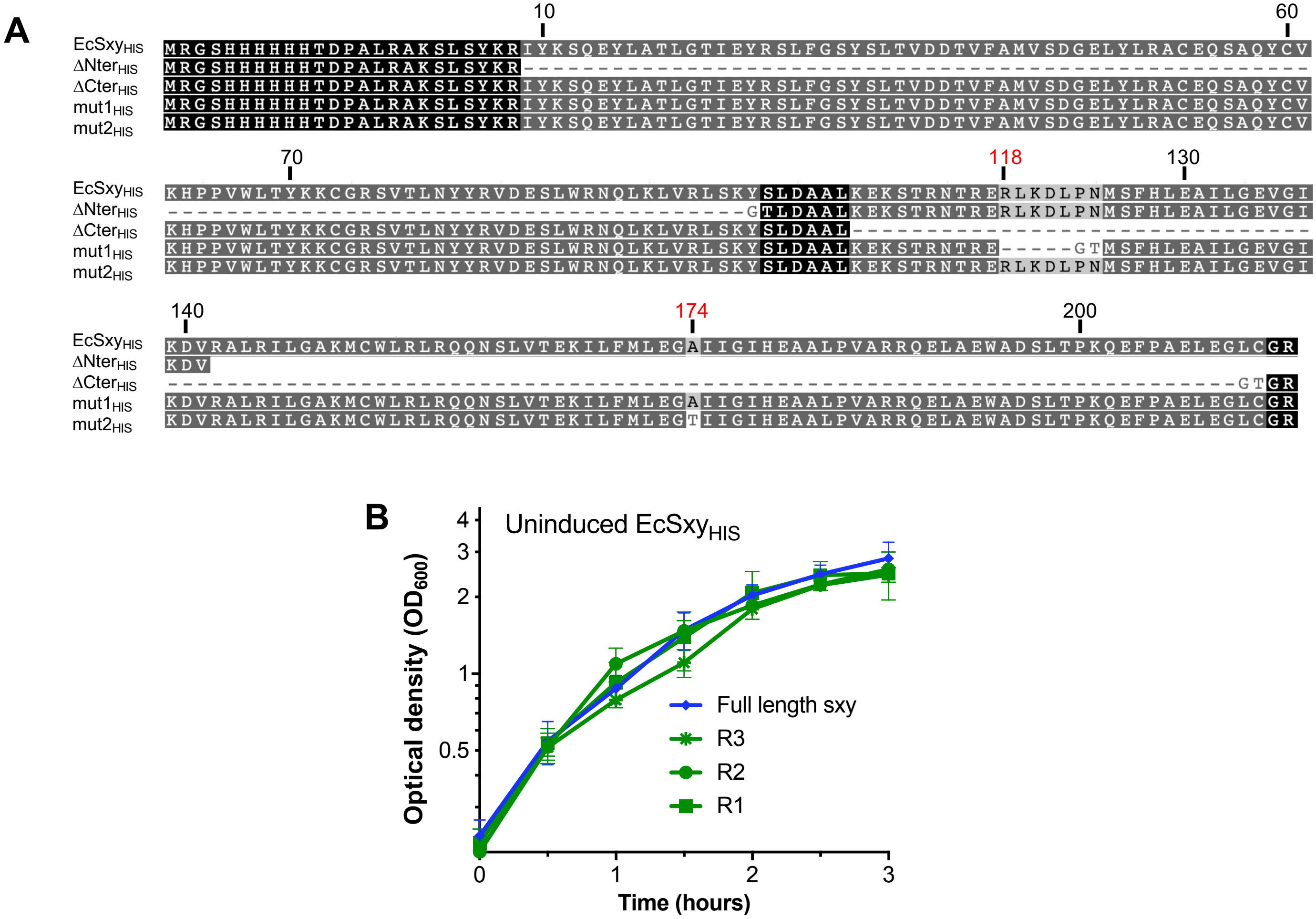

### Supplementary Materials

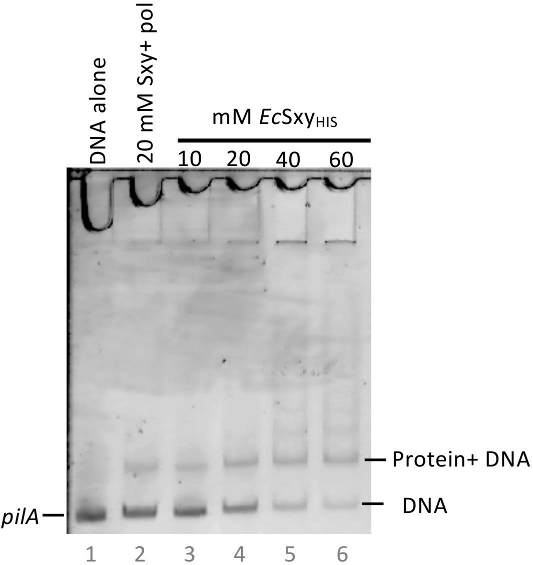


**Bandshift analysis for *Ec*Sxy$_{HIS}$ and *Ec*CRP$_{HIS}$ proteins binding to *H. influenzae pilA* DNA.**

### Supplementary Materials

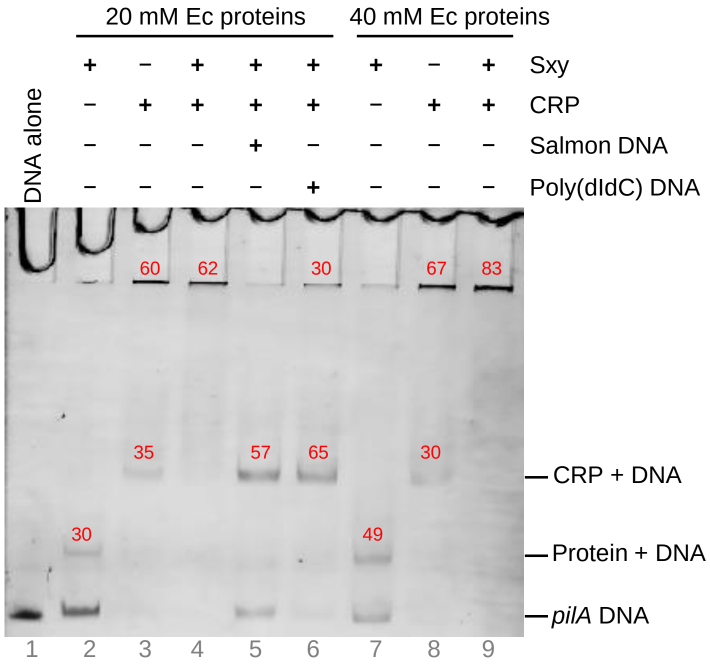
