## Supplementary Materials for "Finding reliable phenotypes and detecting artefacts among *in vivo* and *in vitro* assays to characterize the refractory transcriptional activator Sxy (TfoX) in *Escherichia coli*"

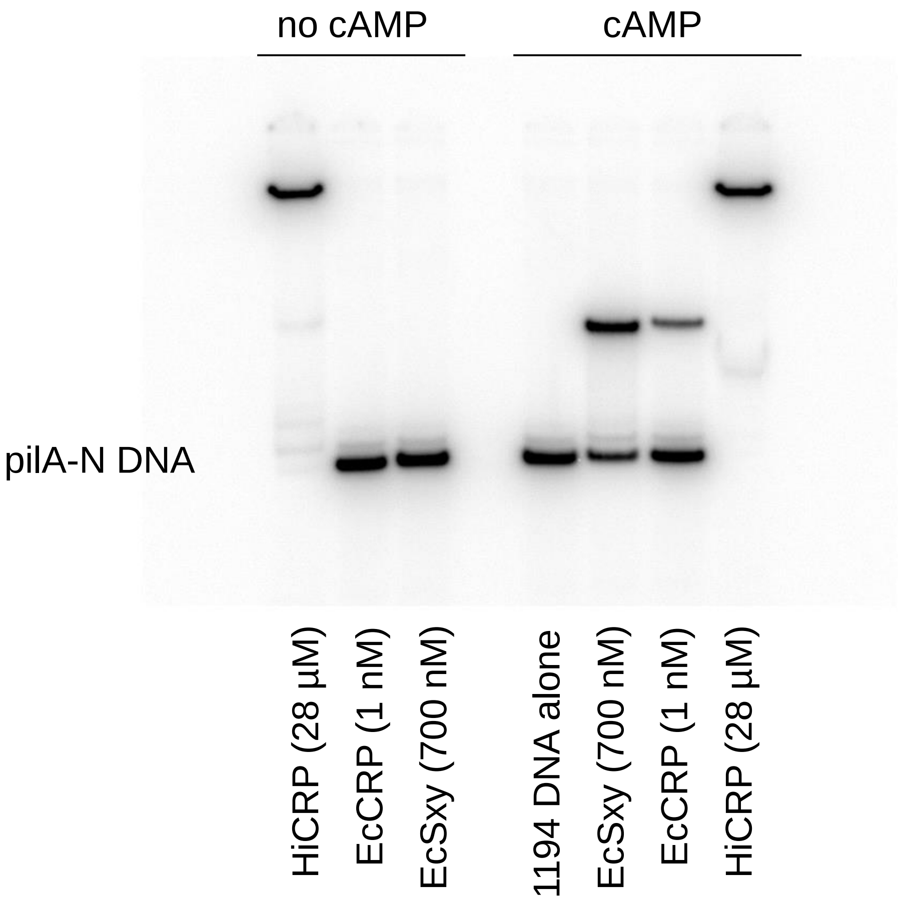


Note: In the absence of cAMP, CRP is known to bind DNA with an affinity ($K_{app}$) around 100 µM (Harman 2001 *Biochimica et Biophysica Acta (BBA) - Protein Structure and Molecular Enzymology* **1547** (1): 1–17). In the above bandshift gel, HiCRP extract is 70,000 x more concentrated than the EcCRP extract. Thus, the complete shifting of bait DNA by HiCRP extract, even in the absence of cAMP, is explained by non-specific DNA binding by CRP at this high concentration. Non-specific binding is consistent with the supershift of the DNA. The supershift could also include contaminating DNA-binding proteins from the HiCRP extract.
