## Supplementary Materials for "Finding reliable phenotypes and detecting artefacts among *in vivo* and *in vitro* assays to characterize the refractory transcriptional activator Sxy (TfoX) in *Escherichia coli*"

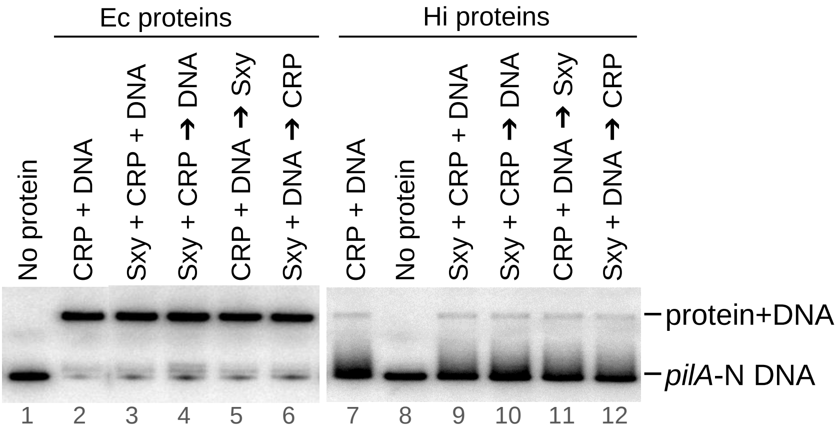


**Bandshift analysis of *H. influenzae* and *E. coli* CRP and Sxy proteins binding to *pilA*-N DNA with sequential incubation of proteins in binding reactions**.

Lanes 1 and 11: DNA alone, Lanes 2 and 12: CRP + DNA, Lanes 3 and 10: Sxy + CRP + DNA mixed together at the same time, Lanes 4 and 9: Sxy + CRP mixed together before addition of DNA, Lanes 5 and 8: CRP + DNA mixed together before addition of Sxy, Lanes 6 and 7: Sxy + DNA mixed together before addition of CRP.  For *E. coli* proteins (lanes 1 to 6), 350 nM Sxy and 4 nM CRP were used. For *H. influenzae* proteins (lanes 7 to 12), 700 nM Sxy and 400 nM CRP were used.
