## Supplementary Materials for "Finding reliable phenotypes and detecting artefacts among *in vivo* and *in vitro* assays to characterize the refractory transcriptional activator Sxy (TfoX) in *Escherichia coli*"

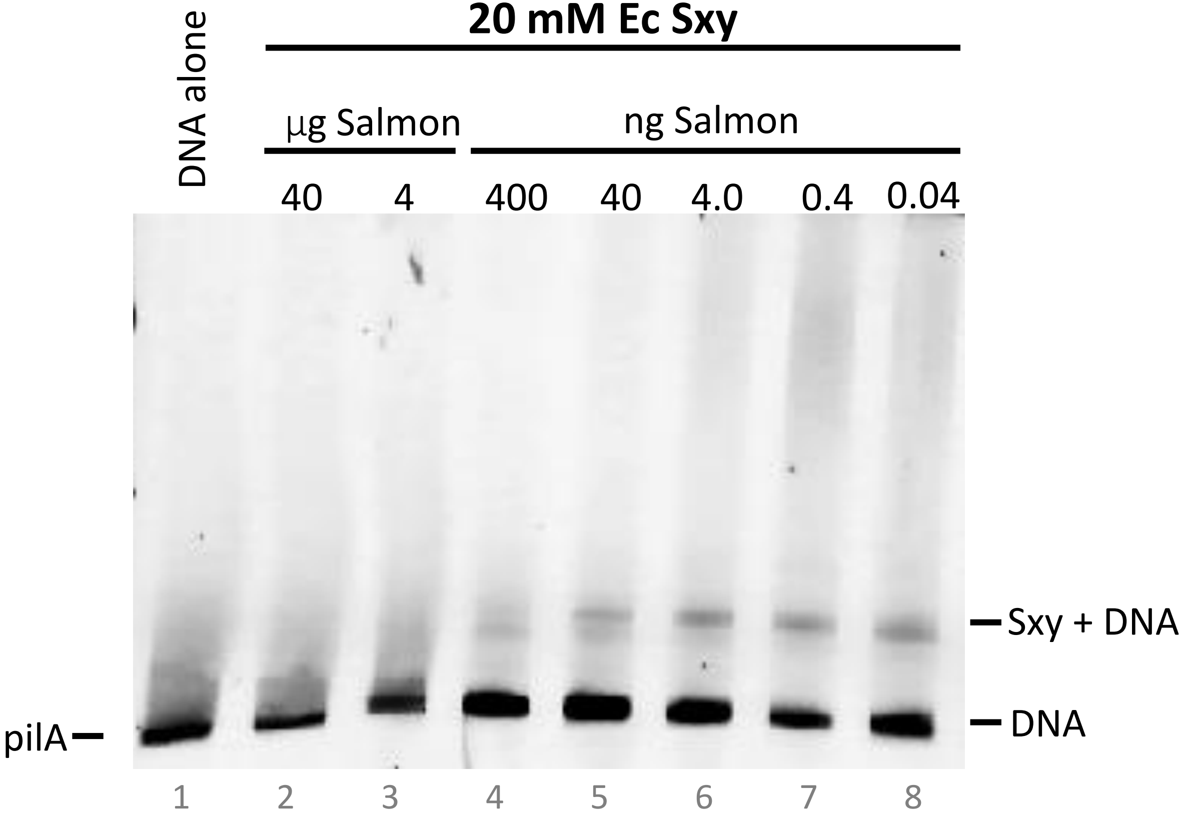


**Bandshift analysis for *E. coli* Sxy proteins binding to *H. influenzae pilA* promoter DNA with titration of Salmon DNA.**
