## Supplementary Materials for "Finding reliable phenotypes and detecting artefacts among *in vivo* and *in vitro* assays to characterize the refractory transcriptional activator Sxy (TfoX) in *Escherichia coli*"

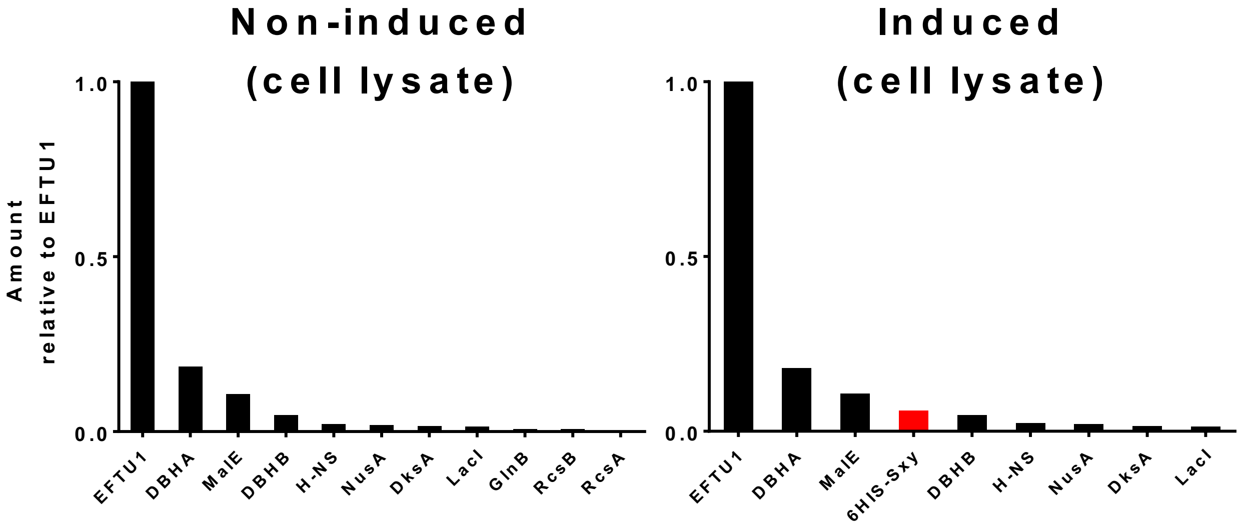


**S7 Fig. Induction and mass spectrometry detection of *Ec*Sxy$_{HIS}$ in whole-cell lysates of *E. coli* Δ*crp*.** Protein abundance is ranked from high to low, with only the most abundant DNA-binding proteins presented in the graphs. Sxy is highlighted in red.
