## Supplementary Materials for "Finding reliable phenotypes and detecting artefacts among *in vivo* and *in vitro* assays to characterize the refractory transcriptional activator Sxy (TfoX) in *Escherichia coli*"

**S1 Table. Carbon substrate fermentation by *E. coli* Δ*crp* cells complemented with cognate or non-cognate CRP and Sxy proteins.** Complementation was conducted with plasmids p*Eccrp*, p*Hicrp,* p*Hisxy*, or p*Ecsxy*. Fermentation was assessed on Difco MacConkey indicator plates containing 1 % of the indicated carbon source.

| | Strain | CRP-independent carbon sources | | CRP-dependent carbon sources | | | |
|---|---|---|---|---|---|---|---|
| | | Fructose | Galactose | Maltose | Glycerol | Mannitol | Xylose |
| 1 | WT | + | + | + | + | + | + |
| 2 | Δ*crp* | + | + | - | - | - | - |
| 3 | Δ*crp* + ***Eccrp*** | + | + | + | + | + | + |
| 4 | Δ*crp* + ***Hicrp*** | + | + | - | - | - | + |
| 5 | Δ*crp* + ***Hicrp*** + ***Hisxy*** | + | + | - | - | - | + |
| 6 | Δ*crp* + ***Hicrp*** + ***Ecsxy*** | + | + | - | - | - | + |
| 7 | Δ*crp* + ***Eccrp*** + ***Hisxy*** | + | + | + | + | + | + |
| 8 | Δ*crp* + ***Eccrp*** + ***Ecsxy*** | + | + | + | + | + | + |

**Supporting text: Sxy does not enhance CRP function at sugar metabolism gene promoters**

CRP is essential for the metabolism of maltose, mannitol, xylose and glycerol, but not fructose or galactose (Table1, rows 1 and 2). Exogenous expression of *Ec*CRP from plasmid p*Eccrp* fully restored wildtype metabolism in an Δ*crp* mutant (S1 Table 1, row 3). In contrast, complementation by p*Hicrp* was only partial: *Hi*CRP restored only xylose fermentation (S1 Table 1, row 4), consistent with an inability for *Hi*CRP to activate all *E. coli* CRP-N promoters because of *Hi*CRP's lower affinity for weak CRP-N sites [1]. This raised the possibility that Sxy stabilization of CRP-DNA interactions could restore maltose, glycerol, and mannitol fermentation. Although co-expression of *Hi*CRP and *Hi*Sxy restores CRP-S promoter function in *E. coli* [2], the same co-expression did not restore sugar fermentation in our phenotypic assays (S1 Table 1, rows 5 and 6). *Ec*CRP activity at CRP-N promoters was unaffected by expression of either *Ec*Sxy or *Hi*Sxy (S1 Table 1, rows 7 and 8). These results suggest that Sxy cannot enhance the binding of CRP to weak CRP-N sites sufficiently to activate transcription.

**S2 Table. List of primer sequences.**

| — | Primer | Sequence (5' -3') | Source |
|---|---|---|---|
| **Mutants** | EcsxyF_Nter | CTCGATGCAGCGCTGAA | This study |
| | EcsxyR_Nter | CCGCTTATAGGAGAGGCTTT | This study |
| | EcsxyF_Cter | GGCCGCTAAGGGTCGACCT | This study |
| | EcsxyR_Cter | CAGCGCTGCATCGAGAGAA | This study |
| | EcsxyF_Mut1 | ATGTCTTTTCATCTGGAAGCG | This study |
| | EcsxyR_Mut1 | TTCCCGGGTATTGCGC | This study |
| | EcsxyF_Mut2 | GCTTGAAGGTAATATTATCGGCAT | This study |
| | EcsxtR_Mut2 | CATGAATGCCGATAATATTACCTT | This study |
| ***sxy* truncati ons** | F1 ec.sxytox1.F | TAAGACCTGCAGCCAAGCTTA | This study |
| | R1 ec.sxytox2.R | TTCCCGGGTATTGCGCG | This study |
| | R2 ec.sxytox3.R | TATACGTAACGCCCGTACAT | This study |
| | R3 ec.sxytox4.R | ACCTTCAAGCATAAACAG | This study |
| **Δ*crp*** | Ec.crp.KO.F | AGCGGCGTTATCTGGCTCTGGAGAAAGCTT ATAACAGAGGGTGTGTAGGCTGGAGGCTGC TTC | This study |
| | Ec.crp.KO.R | GAAACAAAATGGCGCGCTACCAGGTAACGC GCCACTCCGATATGAATATCCTCCTTAG | This study |
| **qPCR** | mglBF | GTCCAGCATTCC- GGTGTTTGG | [3] |
| | mglBR | CGCCTGGTTGTTAGCATCGT | [3] |
| | ppdDF | CGTTTTCGCTAATAGTTGACAG | [3] |
| | ppdDR | AGATTCCGAGGTTTTTTATTTC | [3] |
| | hns-For | CACTTGAAACGCTGGAAGAAATG | This study |
| | hns-Rev | GAGTGCGCTCTTCTTCAACTTCA | This study |
| | pilA.RT-For | CATCGCGGGCGGATAAT | This study |
| | pilA.RT2.Rev | GGAAAGCAGATTCCGAGGTT | This study |
| | comM.qPCR. For | GTTGTTCCCTTCCTCGTAGTT | This study |
| | comM.qPCR.Rev | GTGGCTGATTCCTGTTTATTTGT | This study |
| **DNA baits** | M13F.PmglB.For | TGTAAAACGACGGCCAGT CGTGCCAGCCACTGTTT | This study |
| | M13F.PmglB.Rev | TGTAAAACGACGGCCAGT CGTGCCAGCCACTGTTT | This study |
| | M13F.PppdA.For | TGTAAAACGACGGCCAGT GACTTTGAGATTGAAGGCTACG | This study |
| | M13F.PppdA.Rev | CAGGAAACAGCTATGACCATG CAGTATGGAGCGAGGAGAAA | This study |
| | M13F.PppdD.For | TGTAAAACGACGGCCAGT AACGGGCGTGGACTTTATC | This study |
| | M13F.PppdD.Rev | CAGGAAACAGCTATGACCATG GATTTGGTTGGCGCTACTTTG | This study |
